# Patterns of structural variation in human cancer

**DOI:** 10.1101/181339

**Authors:** Yilong Li, Nicola D Roberts, Joachim Weischenfeldt, Jeremiah A Wala, Ofer Shapira, Steven E. Schumacher, Ekta Khurana, Jan Korbel, Marcin Imielinski, Rameen Beroukhim, Peter J Campbell, on behalf of the PCAWG-Structural Variation Working Group, and the PCAWG Network

## Abstract

A key mutational process in cancer is structural variation, in which rearrangements delete, amplify or reorder genomic segments ranging in size from kilobases to whole chromosomes. We developed methods to group, classify and describe structural variants, applied to >2,500 cancer genomes. Nine signatures of structural variation emerged. Deletions have trimodal size distribution; assort unevenly across tumour types and patients; enrich in late-replicating regions; and correlate with inversions. Tandem duplications also have trimodal size distribution, but enrich in early-replicating regions, as do unbalanced translocations. Replication-based mechanisms of rearrangement generate varied chromosomal structures with low-level copy number gains and frequent inverted rearrangements. One prominent structure consists of 1-7 templates copied from distinct regions of the genome strung together within one locus. Such ‘cycles of templated insertions’ correlate with tandem duplications, frequently activating the telomerase gene, *TERT,* in liver cancer. Cancers access many rearrangement processes, flexibly sculpting the genome to maximise oncogenic potential.

## INTRODUCTION

Mutations arising in somatic cells are the driving force of cancer development. An especially important class of somatic mutation is structural variation, in which genomic rearrangement acts to amplify, delete or reorder chromosomal material at scales ranging from single genes to entire chromosomes. A structural variant (SV) manifests as a join (or ‘breakpoint junction’) linking two segments of the genome not normally juxtaposed, typically associated with an increase or decrease in the copy number of adjacent genomic regions. Simple classes of SV, such as deletion, tandem duplication or unbalanced translocation, have characteristic patterns of breakpoints and copy number change **(Extended Figure 1).** Some cancers are associated with large numbers of such simple events – breast and ovarian cancers can have hundreds of tandem duplications *^1,2^*, for example, while some leukaemias have high proportions of deletions^3^.

A range of more complex structural variation processes have recently been described, often characterised by spatial clustering of several breakpoints **(Extended Figure 1).** Breakage-fusion-bridge events, first described in maize 70 years ago^4^, result from repeated cycles of DNA breakage, end-to-end chromosome fusions leading to mitotic bridges and further DNA breakage. In cancer genomes, these manifest as several proximate, inverted breakpoint junctions with substantial copy number amplification, which we call fold-backs^5–7^. Chromothripsis^8^, in which chromosome shattering and random rejoining occur in a single catastrophic event^9,10^, leads to a pattern of oscillating copy number changes and localised clustering of breakpoints^11^. Chromoplexy^12,13^, particularly frequent in prostate cancers, results from simultaneous dsDNA breaks in several chromosomes that are rejoined incorrectly, leading to balanced chains of rearrangements. Finally, in the germline, replication-based mechanisms of rearrangement can induce small duplications and triplications, thought to arise from stalling of the replication fork leading to template switching^14–16^ – whether this process is also active in cancer remains unknown.

Despite this progress, our knowledge of structural variation in human cancers remains incomplete. Methods for formally reconstructing a cancer genome’s evolution from its set of breakpoints and copy number changes have been developed^17^, but are difficult to apply in practice due to missing SV calls and uncertainty in copy number estimates. Here, we develop a pragmatic system for classifying and annotating SVs (Figure 1A-C; **Supplementary Methods),** and apply it to whole cancer genomes, collected and analysed under the auspices of the PCAWG consortium.

**Figure 1.**
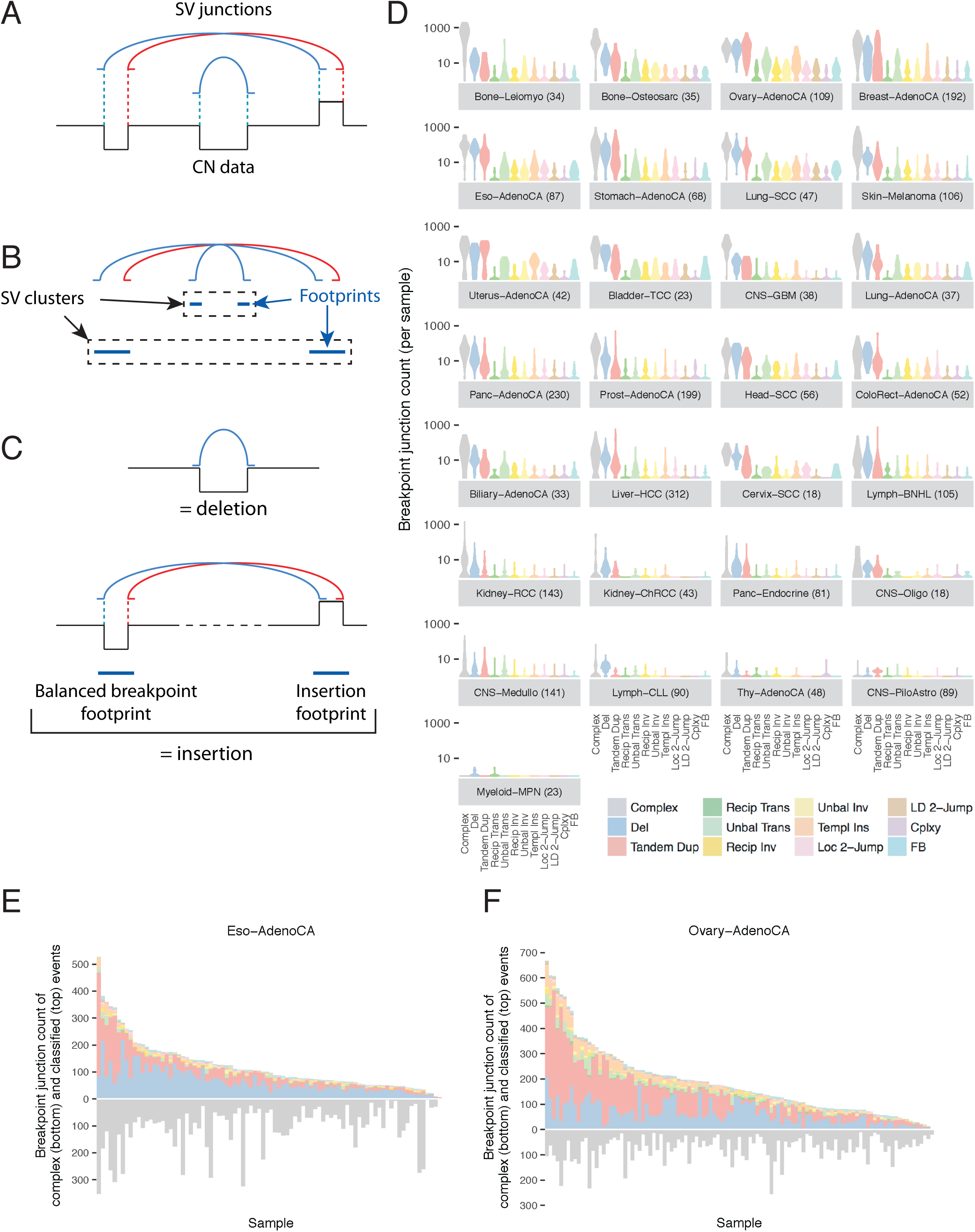
Classification of structural variants in cancer genomes. (A) Structural variants are grouped into ‘clusters’, based on statistical assessment of geographical proximity that incorporates information on each tumour’s number and size distribution of rearrangements. (B) Clusters, that often span large genomic regions or several chromosomes, are then separated into constituent local ‘footprints’ (C) Footprints are then classified into mechanistic categories based on copy number changes and rearrangement orientation. Simple categories include ‘deletions’, ‘tandem duplications’, ‘reciprocal inversions’, ‘fold-back inversions’, ‘unbalanced translocations’ and ‘reciprocal translocations’. More complex categories include ‘chromoplexy’ and a range of structures discussed in the manuscript. (D) Violin plots of density of classified SV categories across patients within each histology group, with number of samples indicated in parentheses. (E) Per-sample counts of complex (lower) and classified (upper) SV breakpoint junctions for oesophageal adenocarcinoma. (F) Per-sample counts of complex (lower) and classified (upper) SV breakpoint junctions for ovarian adenocarcinoma.

### Simple rearrangement classes

Since rearrangements from a given cancer are often highly clustered, we group rearrangements into ‘clusters’ based on the proximity of adjacent breakpoints, the overall number of events in that genome and size distribution of those events (Figure 1A). Given that these clustered rearrangements can bring together distant regions of the genome, we then divide such clusters into local ‘footprints’ (Figure 1B). Isolated breakpoints, not clustered into complex footprints, are classified into well-described categories of simple SV based on the orientation of chromosomal segments and copy number changes (Figure 1C). To evaluate clusters of rearrangements, we built a library of all possible genomic configurations that could be generated from sequential application of simple SV types, to a depth of five rearrangements. For the smaller clusters in the cancer genomes, we could then compare the observed patterns to this library to determine how they might have evolved.

This methodology has the advantage that breakpoint junctions are classified according to the wider genomic context in which they occur. This means that true deletions, say, can be separated from breakpoint junctions that happen to have a deletion-type orientation but arise within, for instance, a chromothripsis event of markedly different mechanism and properties. More than half the breakpoint junctions observed in this dataset arise within complex clusters – removing these from catalogues of true deletions, tandem duplications and inversions enables more precise description of their size ranges, genome-wide distribution and signatures. The methodology also enables us to explore novel patterns of complex rearrangement.

We studied breakpoints identified from aberrantly mapping reads in paired-end sequencing data^18^ across 2,559 whole cancer genomes, of which 2,429 had informative SVs. Complex clusters of rearrangement accounted for the greatest number of breakpoints across most tumour types (Figure 1D). Among simple SV categories, deletion was the most common, followed by tandem duplication and unbalanced translocation. Reciprocal translocations and reciprocal inversions were rare events (event classes depicted and defined in **Extended Figure 1).**

There was considerable variability in the overall numbers and distribution of SV classes across tumour types and across patients within a given tumour type (Figure 1E-F; **Extended Figure 2).** For example, oesophageal cancers were characterised by many deletions and a large number of complex clustered rearrangements, even in otherwise quiet genomes. The numbers of rearrangements per patient in ovarian cancer ranged from <10 to several hundred; with some patients carrying high numbers of tandem duplications; others mostly deletions; and a steady frequency of a few unbalanced translocations per patient.

### Complex SVs can be subdivided into constituent local footprints

More than half of the rearrangement junctions in the dataset did not derive from simple SV classes such as deletion or tandem duplication. To explore potential mechanisms of rearrangement underlying more complex events, we analysed the sets of local footprints that comprise individual constituents of complex clusters, with a focus on recurrent footprint patterns (also discussed in considerably greater detail in **Supplementary Results).** The simplest and most recurrent local footprint in the cohort was of a copy number gain bounded by rearrangements linking either end of the duplicated template to other, distant regions of the genome – these we call ‘templated insertions’ (Figure 2A(i)). In such cases, the two rearrangement junctions are likely to be ‘phased’ (occur on the same derivative chromosome haplotype). A second footprint associated with rearrangements linking to distant regions of the genome is associated with local copy number loss – these we call ‘balanced footprints’ (Figure 2A(ii)). Such events are ‘unphased’, since the respective orientation of the two local rearrangements means that they cannot both reside on the same derivative chromosome. A reciprocal translocation is an example of two balanced footprints linked together, while chromoplexy comprises multiple interlinked balanced footprints^12,13^.

**Figure 2.**
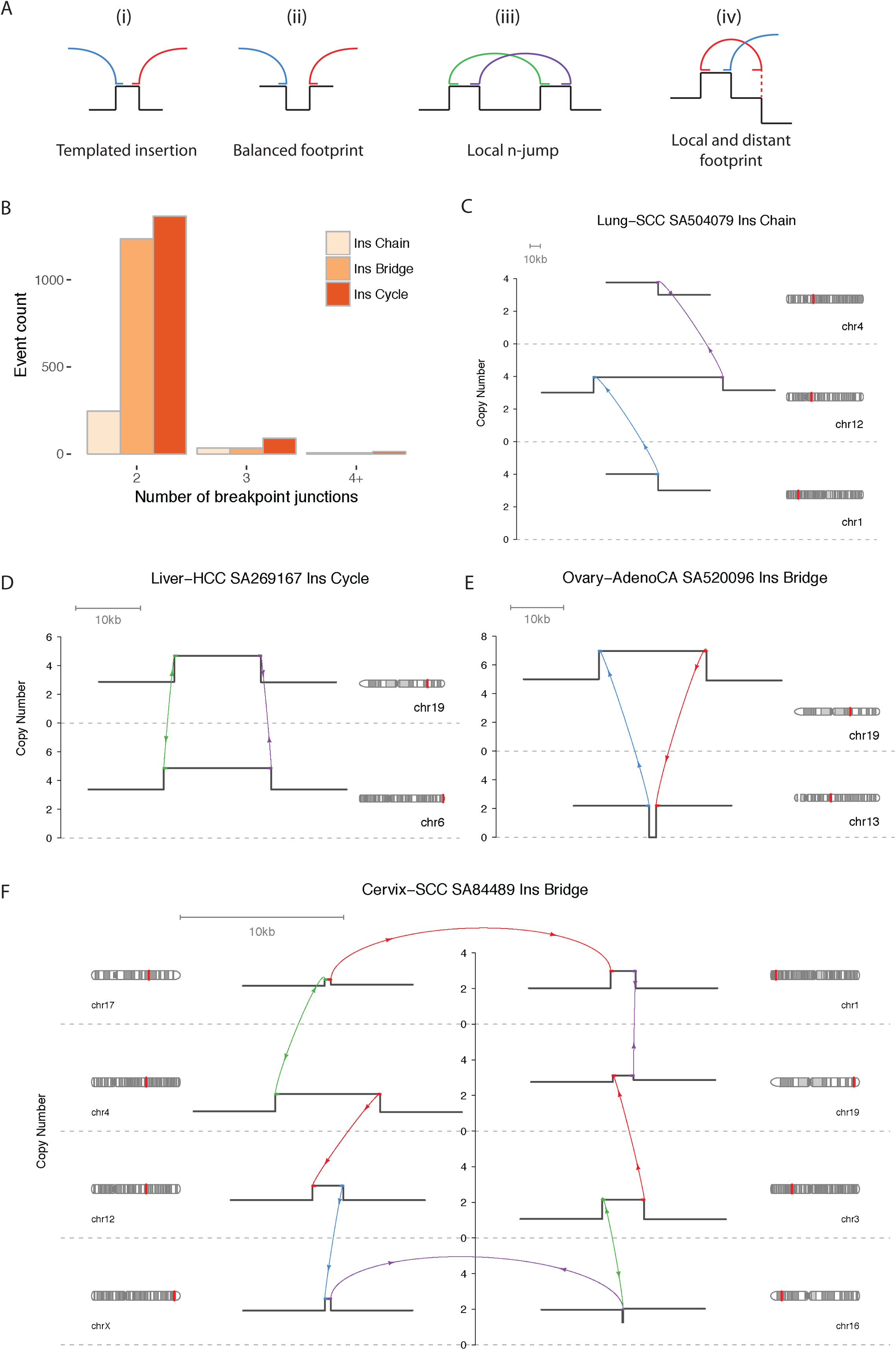
Chains, cycles and bridges of templated insertions. (A) Four broad classes of local footprint structures. Shown in black are relative copy number structures, with the coloured arcs representing rearrangement junctions. At the base of each arc is a flat line showing the orientation of the rearrangement join. Arcs that have only one end on the local footprint represent rearrangements linking to distant regions of the genome. (B) Counts of templated insertion chain, bridge and cycle events with two, three, or more breakpoint junctions, tallied across the whole cohort. (C-E) Examples of typical chain, cycle and bridge events respectively. The estimated copy number segmentation is drawn in black, and breakpoint junctions are indicated by coloured arrows. (F) The longest templated insertion event was an insertion bridge with seven templated insertions strung together and dropped into a region on chromosome 16.

In addition to footprints connected to distant regions of the genome, we found a number of recurrent footprint patterns characterised by several, purely local rearrangements, which we call ‘local n-jumps’ with n denoting the number of rearrangement joins (Figure 2A(iii)). As might be anticipated, there were also recurrent footprints that combined local and distant rearrangements (Figure 2A(iv)). In addition to the recurrent footprint patterns described in the following sections, there were also many local footprint patterns that were observed only once in the cohort. These typically comprised many rearrangements, with their unique footprint presumably arising from the combinatorial explosion of possible structures as numbers of rearrangements increase.

### Chains, cycles and bridges of templated insertions

A particularly striking class of linked footprints consists of several templated insertions from across the genome strung together in a single haplotype (Figure 2B-F). These fall into three basic categories defined by whether or not the string of inserted segments returns to the original locus: those that do not return we term *chains* of templated insertions; and those that do return are either *bridges* (leaving a gap on the host chromosome) or *cycles* (re-replicating a segment on the host).

Cycles and bridges of templated insertions involve segments with the same absolute gain in copy number relative to their source chromosomes, linked so that a path moves once through each segment before returning to the original chromosome of departure. We observed 1467 cycles where the point of return is behind the point of departure – in these cases there is a duplication on the host chromosome and the identity of the host chromosome is not easily determined. We also observed 1275 bridge events where the point of return is after the point of departure, generating a deletion on the host chromosome (Figure 2D-F; Extended Figure 3).

In *chains of templated insertions,* the string of genomic segments does not return to the locus of departure (Figure 2C, **Extended Figure 3),** but is similarly associated with copy number gains at each templated segment. There were 285 instances of such chains in the dataset. One of the commoner manifestations is of an unbalanced translocation in which multiple templated insertions are strung together between the two aberrantly joined chromosomes.

Most templated insertion events involve just two breakpoint junctions (generating one amplified insert in chains and bridges, and two in cycles), but this can extend to three, four or more linked rearrangements (Figure 2B). The longest such event, from a cervical squamous cell cancer, had seven templated insertions strung together on an eighth host chromosome (Figure 2F; other long examples in **Extended Figure 3).**

### Local *n*-jumps

There were a number of recurrent footprints in the dataset which solely contained rearrangements confined to one genomic region. Of those comprising two local rearrangements, some had straightforward explanations such as nested or adjacent tandem duplications. Many, however, did not have a trivial explanation (Figure 3A). These included a duplication-inverted-triplication-duplication structure that has been observed in germline SVs^16^ (349 instances); a structure of two duplications linked by inverted rearrangements (531 instances); and structures of copy number loss plus nearby duplication linked by inverted rearrangements (472 instances). These patterns all had theoretical solutions recapitulating the observed copy number profiles with breakpoints phased to a single haplotype (Figure 3A), but these configurations could not plausibly be generated by the sequential operation of simple SVs.

**Figure 3.**
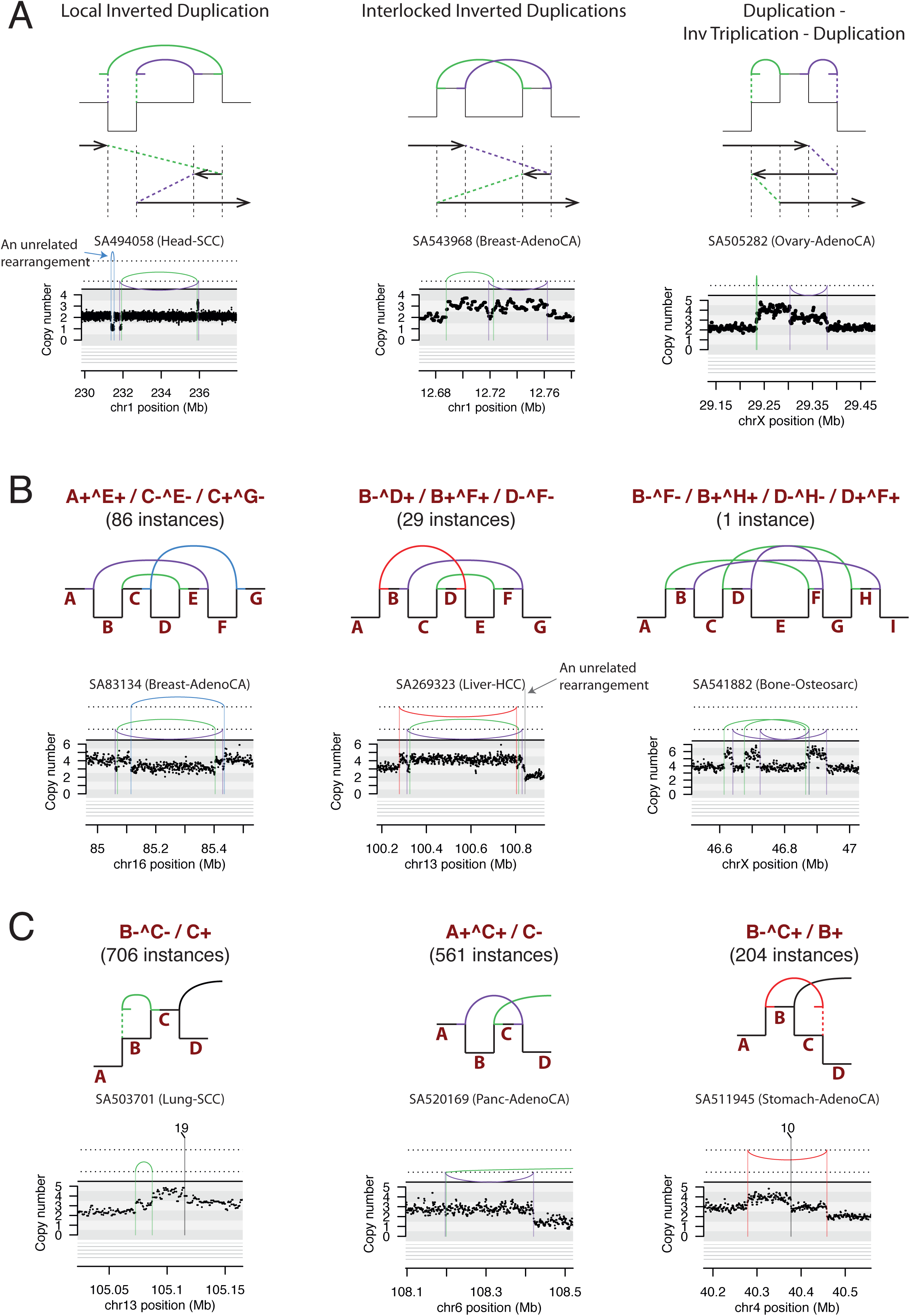
Examples of clusters of 2-5 rearrangements seen in human cancers. (A) Structures created by two local rearrangements that cannot easily be explained by simple SV categories. The upper third of each example shows the copy number and SV patterns observed; the middle third shows one phased genomic sequence that would recreate the observed pattern; and the lower third shows an example observed in a given tumour. (B) Structures created by 3-4 local rearrangements that cannot easily be explained by simple SV categories. The notation for describing the structures uses capital letters (A, B, C…) to denote genomic segments demarcated by rearrangement junctions. The left and right ends of these segments are denoted by + and – respectively. A rearrangement joining two segment ends within the footprint is indicated using ^ (for example, A+^C-denotes a rearrangement joining the right end of segment A to the left end of segment C). Rearrangements that reach from the local footprint elsewhere in the genome are represented by just the segment end separated from other rearrangements by /. (C) Structures created by one local rearrangement and one rearrangement reaching elsewhere in the genome.

To exemplify our reasoning, consider the rearrangement structure of two duplications linked by inverted breakpoint junctions (Figure 2A(iii)). Using our genomic configuration library of all possible sequential SV combinations, we can define four possible routes to this structure **(Extended Figure 4).** The first is an episomal circle comprising the two amplified segments, but this is an unlikely mechanism because the absence of a centromere leads to random episome segregation at mitosis and instability of copy number per cell^19^. In contrast, most of our examples were at stable, integer copy numbers. The second possible explanation is two foldback rearrangements on different copies of the chromosome, but this cannot explain all instances because linked, inverted duplications were sometimes found in tumours with only one copy of that chromosome. Thirdly, two unbalanced translocations between sister or homologous chromosomes, while formally possible, is unlikely because the average copy number on each side of the event for affected chromosomes is no lower than the rest of the genome on average. Finally, a tandem duplication followed by a reciprocal inversion and then a deletion could create the observed structure, but, if so, we would expect to see many more instances of the intermediate stage of tandem duplication with inversion. In fact, the linked, inverted duplication structure is far more common in this cohort (531 instances) than an inversion within a tandem duplication (33 instances).

Beyond clusters of two rearrangements, we also found examples involving three, four or more rearrangements confined to one genomic locale (Figure 3B). These could all be phased to a single haplotype, with breakpoints tightly grouped. As above, although there are routes to constructing these clusters through sequential operation of simple SVs, these solutions require implausible machinations of sister or homologous chromosomes. Instead, we will reason below that local *n*-jump structures are most likely the result of a replication-based mechanism of rearrangement.

### Footprints with local and distant rearrangements

Many local footprints combined local jumps with rearrangements reaching into one or more distant regions of the genome (Figure 3C). Simple examples of these events include unbalanced translocation or large deletion with a locally-derived fragment inserted at the breakpoint, but there was also an extensive range of more complex patterns **(Extended Figure 5).** In some cases, the source of the inserted fragment was distal to the major break, and the SV could feasibly result from several concurrent DNA breaks in close spatial proximity with capture of a short DNA fragment during repair. In other cases, the origin of the inserted fragment was proximal to the major break. Inserted fragments from both distal and proximal loci were typically associated with a gain in copy number. This is difficult to explain by a fragmentation and ligation mechanism because the copy number gain implies the inserted segment was a duplicate of the original template, rather than a separated fragment redistributed from its original locus.

Curiously, a comparison of footprints linked together through distant rearrangements revealed strong connectivity of footprints with the same or similar local structure, often enriched 10-fold or more than expected by chance (see **‘Footprint connectivity analysis’, Supplementary Results).** The reasons for this are unclear, but may reflect innate structural symmetry introduced either through the generation or resolution of rearrangements, or through the repeated action of a mechanism imparting consistent structural motifs.

### Replication-based mechanism of structural variation

The diverse structural variation patterns described above **(Figures 2-3)** share important morphological features: (1) genomic configurations that can be phased to a single haplotype; (2) low-level gains in copy number, especially duplications and triplications; (3) a high frequency of inverted rearrangements in addition to noninverted rearrangements; (4) occurrence on a chromosome background with similar average copy number to the tumour overall; and (5) tight proximity of breakpoints within the local footprint, typically <1Mb. We can be confident that these structures do not result from sequential operation of simple SVs, as reasoned above. In particular, the prevalence of inverted breakpoint junctions and local copy number gains is difficult to recreate using simple rearrangements without recourse to multiple unbalanced recombination events across sister or homologous chromosomes, an unlikely solution.If these events cannot be satisfactorily explained by sequential simple rearrangements, another possible explanation is a mechanism involving simultaneous DNA breakage and aberrant ligation of fragments. Chromothripsis, chromoplexy and breakage-fusion-bridge cycles are examples of such processes occurring in a single crisis during cancer evolution. However, the patterns described above do not fit the known properties of these mechanisms either (for more detailed analysis, see **Supplementary Results).** The regional shattering of classical chromothripsis typically generates chromosomal loss^8,20^, as opposed to the copy number gains described here. Although chromothripsis in association with copy number gain has been described, either preceding local genomic amplification^11,21^ or after whole chromosome duplication or breakage-fusion-bridge cycles^7,22,23^, the resulting copy number and rearrangement patterns are different to those described here. Chromoplexy, in which chromosome breaks lead to balanced interchange at multiple breakpoint junctions^12,13^, typically generates unphased solutions, whereas the events described here can be phased to a single haplotype. Repeated breakage-fusion-bridge cycles tend to cause high-level copy number gains associated with inverted, fold-back rearrangements^5,6,24^, unlike the structures reported here.

Instead, we believe many of these locally confined SV clusters with low-level copy number gains are generated in a single event by a replication-based rearrangement mechanism. A replication mechanism has previously been proposed for the duplication-inverted-triplication-duplication structure and other events seen in the germline^14-16^. Under this hypothesis^14-16^, stalled replication forks or other DNA lesions cause the DNA polymerase to switch templates and continue replication in the new location. The new replication fork is itself prone to stalling and further template switching, generating increasingly complex rearrangement structures. Eventually the DNA polymerase switches back to the original template strand, in the correct orientation, and DNA replication can continue unimpeded. If all the template switches are local, this process would explain all observed characteristics in **Figure 3A-B:** the tight clustering of breakpoints, the phased solutions, the random orientation of breakpoint junctions, the frequency of copy number gains, and the average copy number of the background chromosome. If the template switch includes a local jump and a jump to another chromosome, it would generate an unbalanced translocation with an intervening inserted fragment **(Figure 3C),** implying that unbalanced translocations may result from a replication-based mechanism. Finally, if all template switches are distant, the replication mechanism would generate chains, cycles and bridges of templated insertions (Figure 2).

### Genomic properties of SVs

The size distribution of tandem duplications showed interesting patterns across tumour types **(Figure 4A).** Stomach and endometrial cancers had predominantly large tandem duplications, mostly >100kb; cervical and colorectal tumours had smaller events in the 10-100kb range; and breast and ovarian cancers had a distinctly bimodal or trimodal distribution. At the individual patient level, however, tumours tended to have a unimodal distribution of tandem duplication size **(Figure 4B).** This suggests that (at least) two separate mechanisms of tandem duplication operate in human cancers, and individual tumours draw predominantly or exclusively from one of these mechanisms. It is known, for example, that breast cancers with *BRCA1* loss have small tandem duplications, whereas those with *BRCA2* loss have a mix of larger tandem duplications and small deletions^25^. Similar themes emerged for deletion size across the cohort **(Extended Figure 6).** Strikingly, the sizes of individual fragments in templated insertion events were also distinctly bimodal, with varying peak heights across tumour types **(Figure 4C).** Curiously, when correlating template sizes within a given event, two patterns emerged – one in which template sizes were closely correlated with one another, and one in which a small (<1kb) template was linked with one of any size **(Extended Figure 6).**

**Figure 4.**
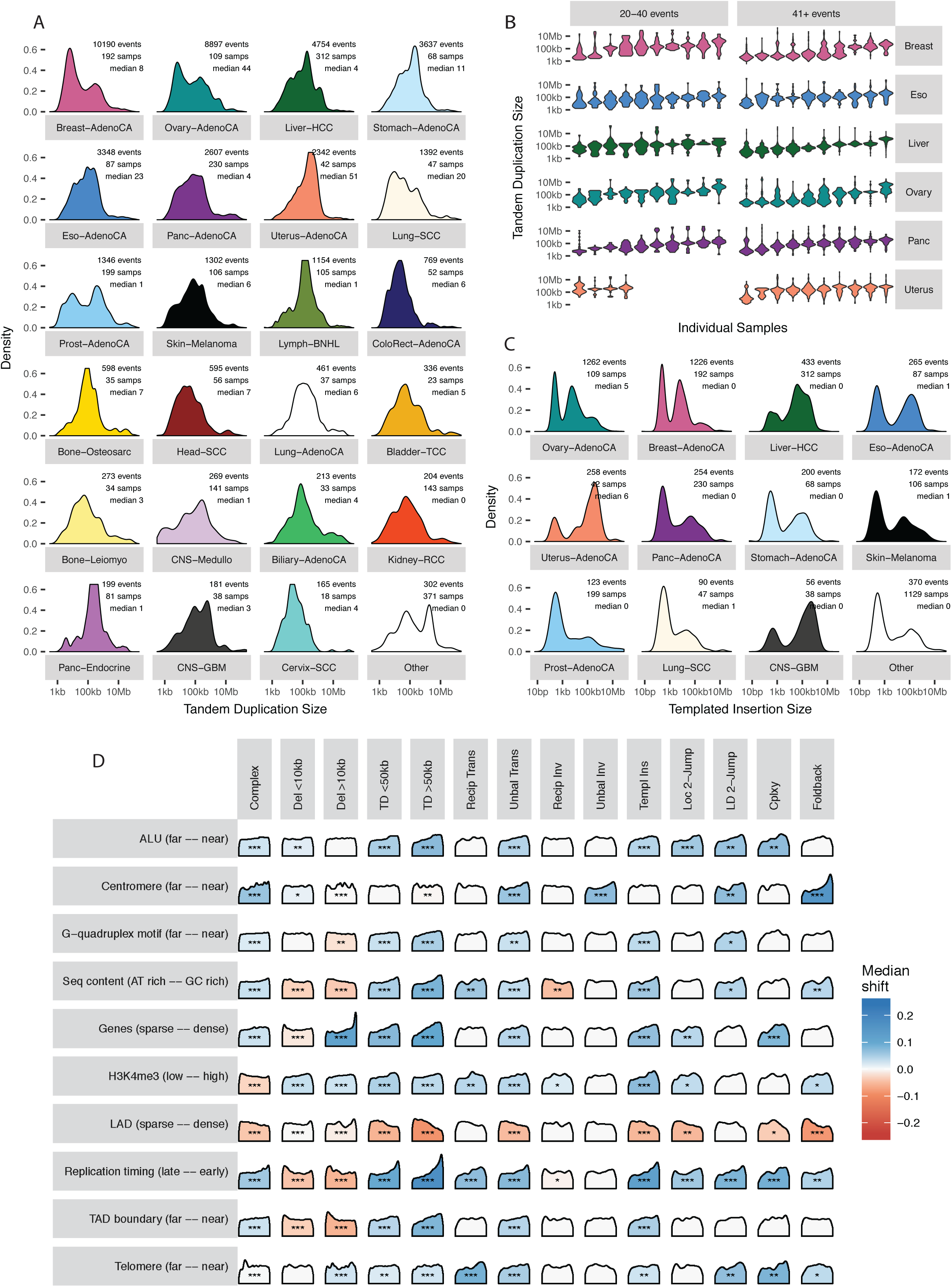
Size distribution and genomic properties of classified SVs. (A) Size distribution of tandem duplications per histology group, annotated with the number of tandem duplications, number of samples, and the median tandem duplication count per sample. (B) Size distribution of tandem duplications per sample, for a subset of individuals at deciles along the range of per-sample median sizes. Samples in the left panel had between 20 and 40 tandem duplications; samples in the right panel had more than 40. (C) Size distribution of segments of templated insertion per histology group, annotated with the number of such inserts, number of samples, and the median templated insert count per sample. (D) Associations between genomic properties (rows) and classes of structural variation (columns). Each density curve represents the quantile distribution of the genomic property values at observed breakpoints compared to random genome positions, with stars indicating significant departure from uniform quantiles: FDR <0.01 *, <0.001 **, and <10^−6^ ***. Cells with significant property associations are shaded by the magnitude of the shift of the median observed quantile above (blue) or below (red) 0.5. The interpretation of each property from left to right is indicated in parentheses.

A number of genomic properties, such as replication timing, transcriptional activity and chromatin state, influence the density of point mutations^26–28^, but what effects they have on SV processes are unclear. From the literature, we compiled a library of the genome-wide distribution of 38 features, including replication timing, GC content, repeat density, gene density and distance to G-quadruplex motifs among others. Replication timing had the strongest association with SV occurrence, with deletions enriched in late replicating regions, and tandem duplications and unbalanced translocations preferentially occurring in early replicating regions **(Figure 4D, Extended Figure 7, Supplementary Figure 1).** Likewise, regions of active chromatin and increased gene density correlated positively with the rate of rearrangement. These correlations are the reverse of those observed for point mutations, which generally have higher density in late replicating, inactive and gene-poor regions of the genome^26-28^.

### Signatures of structural variation

The differences across patients in size distribution of tandem duplication and deletion, together with the widely varying frequency and patterns of SV across tumour types and genome topology, suggest that a number of distinct rearrangement processes can operate in varying combinations to sculpt a cancer genome. These can be separated and described as structural variation ‘signatures’ by fractionating the within-patient cooccurrence patterns between different SV classes, as previously undertaken for point mutations^29^. We applied two statistical methods for signature discovery **(Supplementary Methods),** with overall similar results. We included only those local rearrangement footprints that were recurrent in the cohort. As a result, many of the very complex clusters of rearrangement, such as those arising from chromothripsis^8^ and genomic amplifications, are excluded from the signatures analysis. Overall, we found evidence for nine structural variation signatures in this dataset **(Figure 5; Supplementary Figures 2-3).**

**Figure 5.**
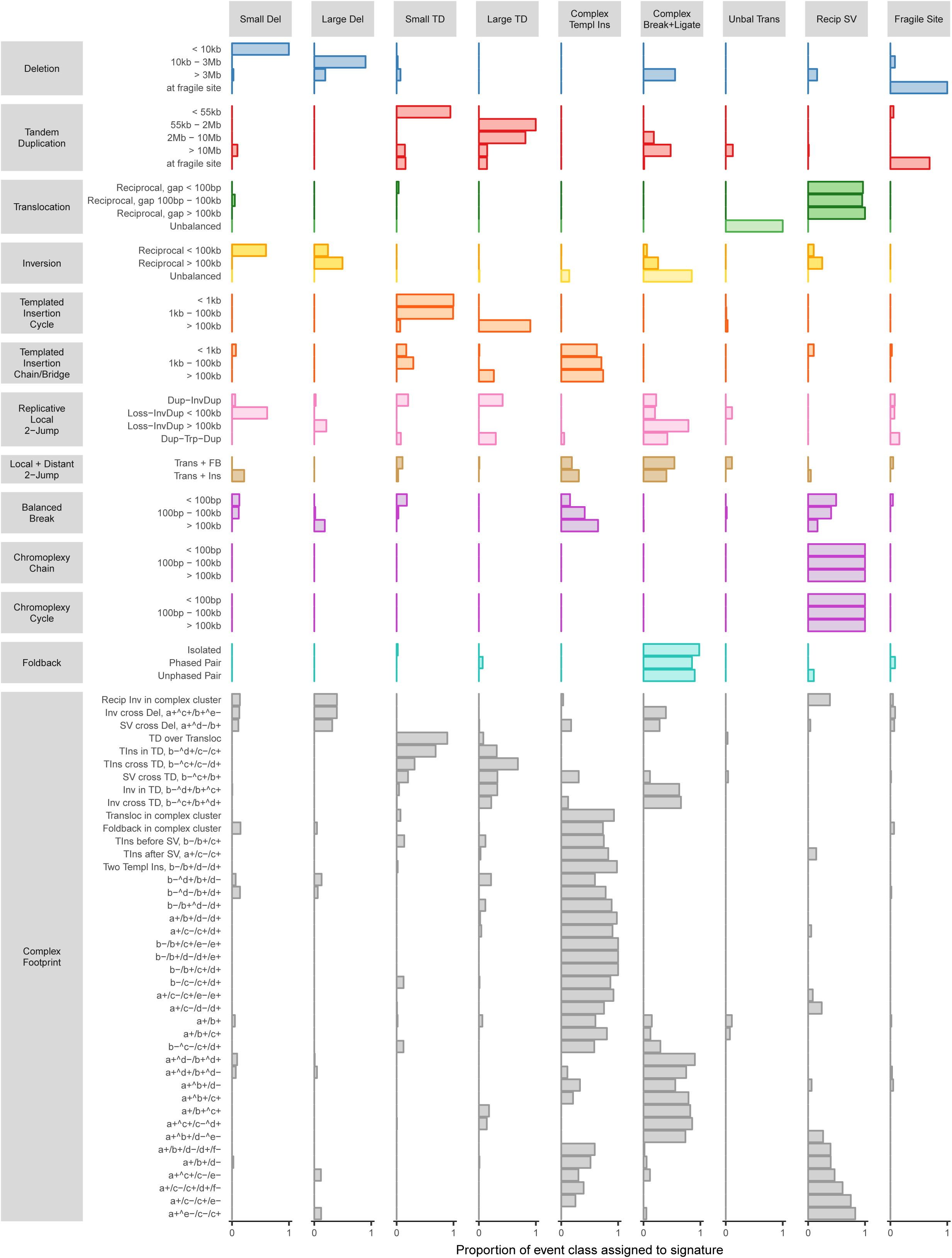
Structural variant signatures in human cancers. The nine structural variation signatures calculated by non-negative matrix factorization. Here we show the estimated proportion of event classes conferred by each signature (columns sum to one).

One signature comprised predominantly small deletions and reciprocal inversions; and a second signature contained the larger deletions and inversions. These results imply that deletions and reciprocal inversions co-occur and are mechanistically linked, probably through a process generating two breaks within a local genomic region. Given the considerably higher frequency of deletion, the intervening DNA segment must generally be lost to the cell, but can occasionally be recaptured in the opposite orientation to generate a reciprocal inversion. The separation of these two signatures by size range presumably reflects distinct mechanisms of DNA breakage and/or repair operating at different ranges of genomic distance.

Two signatures grouped tandem duplications with cycles of templated insertions, again distinguished by size. Small tandem duplications (<55kb) were associated with cycles of short templated insertions (<100kb), while tandem duplications of 55kb-10Mb correlated with cycles of long templated insertions (>100kb). These signatures imply that cycles of templated insertions co-occur and are mechanistically linked with tandem duplication. Small tandem duplications are a feature of *BRCA1* deficiency^25^, suggesting that the homologous recombination pathway is important in suppressing these events in normal cells.

A fifth signature was characterised by deletions and tandem duplications at fragile sites. It has been well recognised that certain ‘fragile’ regions of the genome are prone to deletion^30^, but that they are also enriched for tandem duplications has not been described previously. Interestingly, tandem duplications were more prominent at the edges of the fragile site, whereas deletions concentrated in the centre **(Extended Figure 8A).** The size range of fragile site deletions peaked around 100kb, similar to the larger deletion signature, while the rarer fragile site tandem duplications showed no strong size peak **(Extended Figure 8).** Sites of fragility varied extensively across tumour types **(Extended Figure 8D),** presumably reflecting differences in local chromatin state or replication timing across tissue lineages.

Unbalanced translocations comprised their own signature, suggesting they derive from a distinct rearrangement process in cancer genomes. A further signature linked the fold-back inversions that are a hallmark of breakage-fusion-bridge cycles with large intrachromosomal events spanning many megabases that could assume any orientation. This signature probably reflects ligation repair of distant intrachromosomal dsDNA breaks, as distinct from the locally confined rearrangements seen in the first five signatures. The prominence of breakage-fusion-bridge cycles could indicate a role for telomere dysfunction in this signature^23,31^.

There was a signature of balanced rearrangements, including reciprocal translocations and chromoplexy clusters^12^. This signature probably arises from dsDNA breaks, potentially occurring in interphase, in which both sides of the break are incorrectly repaired through ligation to other, simultaneously broken regions of the genome.

The final signature comprised chains of templated insertions and other complex footprints associated with clusters of local and distant rearrangements. This signature is likely to encompass several underlying mechanisms, including chromothripsis, other breakage-ligation processes and some of the more complex replication-based mechanisms described earlier.

These nine signatures showed considerable heterogeneity in their activity across tumour types and among patients within a given tumour type **(Extended Figure 9).** Tumours of the gastrointestinal tract, including colorectal and oesophageal adenocarcinomas, showed high rates of the fragile site signature. Some ovarian cancers showed predominance of the small tandem duplication signature, others the large tandem duplication signature; endometrial cancers really only sampled the large tandem duplication signature to any degree. Prostate cancer was striking for the prevalence of the chromoplexy signature, as reported previously^12,13^.

### Cycles of templated insertions frequently activate *TERT* in liver cancer

SVs drive tumour development through their effects on cancer genes, whether by altering gene copy number, disrupting tumour suppressor genes, creating fusion genes or juxtaposing the coding sequence of one gene with the regulatory apparatus of another. The events that rearrange these cancer genes therefore reflect the structural variation mechanisms available to the tumour, filtered by selective advantage on the clone. Thus, the *TMPRSS2-ERG* fusion gene of prostate cancer is frequently created by chromoplexy, as this is one of the major SV signatures active in prostate; rearrangements inactivating *PTEN* are frequently (out-of-frame) internal tandem duplications in breast and ovarian cancers, where these signatures are so active; and amplifications of *MDM2* are almost always driven by chromothripsis-like events, in keeping with their predominance in sarcomas **(Extended Figure 10).**

An unexpected observation was that liver cancers had frequent cycles of templated insertions affecting the *TERT* gene **(Figure 6A-B, Supplementary Figures 4-5).** *TERT* promoter point mutations are present in 54% of liver cancers and an additional 5-10% have structural variants activating the gene^32^. Of 30 liver cancer patients with SVs affecting the *TERT* locus, we find that 10 were templated insertion events, mostly cycles. All these events duplicated the entire *TERT* gene and linked it to duplications of whole genes, fragments of genes or regulatory elements from elsewhere in the genome. Unfortunately, with cycles of templated insertions, we cannot unambiguously identify the host chromosome, so cannot distinguish between other regions being inserted into the *TERT* locus versus a copy of the intact *TERT* gene being dropped into a highly active locus elsewhere in the genome. Whichever applies, it is clear that this particular rearrangement process is distinctive for the exquisite precision with which a cancer can copy-and-paste normally disparate functional elements of its genome together without wholesale instability.

**Figure 6.**
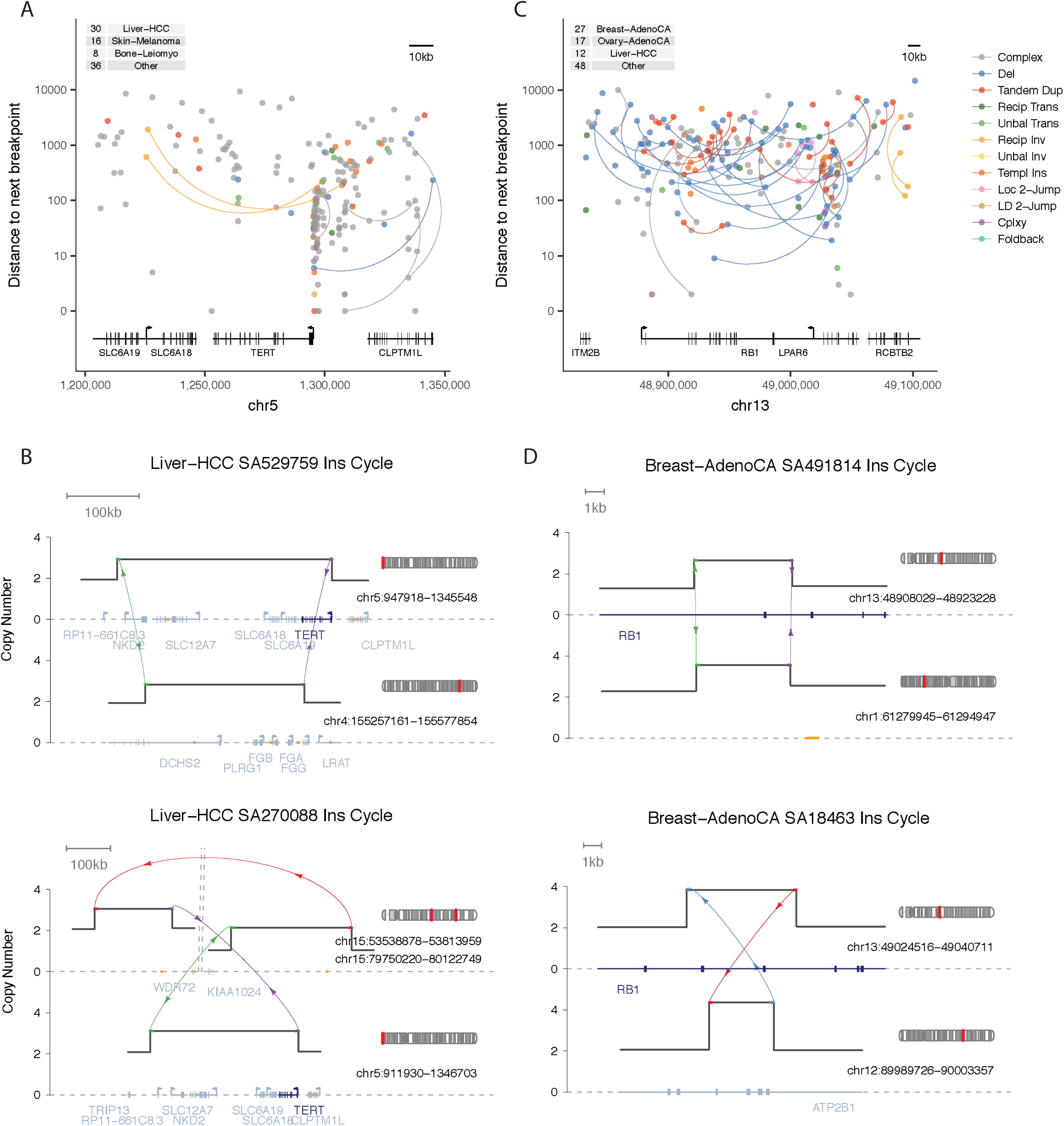
Link between rearrangement mechanism and driver SVs. (A,C) The positions of all SV breakpoints in the *TERT* and *RB1* regions (including 50kb flanks either side), coloured by classification and vertically spaced by the distance to the next breakpoint in the cohort. If the two sides of a breakpoint junction are contained within the plotting window, they are joined by a curved line. The number of samples with a breakpoint in the plotting window is annotated top left. (B,D) Examples of templated insertion cycles affecting (B) the *TERT* locus in liver cancer and (D) the *RB1* locus in breast cancer. Cancer-associated genes are shown in navy; (a subset of) other genes are shown in grey; and enhancer elements are marked in orange.

Replication-based mechanisms were also a frequent source of inactivating SVs in key tumour suppressor genes **(Figure 6C-D, Extended Figure 10B, Supplementary Figures 6-7).** For example, amongst many straightforward deletions, *RB1* was hit by cycles of templated insertions, a templated insertion with deletion and one instance of the linked, inverted duplications detailed above. These events typically generated duplications of internal exons in *RB1* and/or insertions of exons from other genes, all of which presumably rendered a non-functional transcript.

## DISCUSSION

Here, we have taken a constructive approach to defining mechanisms and signatures of structural variation in human cancer genomes. Picking off relatively simple footprints, parsing them into their component events, and comparing across patients enables us to see correlations and contrasts between classes of rearrangement. Already, this has revealed nine structural variation signatures, including some likely to be mediated by replication-based mechanisms. However, we are reaching the limits of such constructive approaches and there remains a large swathe of complex structural variation in cancer genomes that eludes formal characterisation. Undiscovered mechanisms of rearrangement may lurk in the depths of this complexity, or it may be that all can be explained by combinatorial application of known mechanisms.

Genomic instability in cancer, then, is not a single phenomenon. Instead, many different mutational processes can act to restructure the genome and, in doing so, generate a remarkably flexible array of possible structures. Any given person’s tumour draws on a subset of the available processes, shaped by the cell of origin, germline predisposition and other unknown factors. Selection does the rest, promoting the clone that has chanced upon the particular structure that optimises its potential for self-determination.

## ACKNOWLEDGEMENTS

This work was supported by the Wellcome Trust. P.J.C. is a Wellcome Trust Senior Clinical Fellow. The results published here use data generated by the TCGA Research Network: http://cancergenome.nih.gov/. We thank, in particular, members of the Technical Working Group of PCAWG for their assistance in generating the somatic mutation calls that underpin these analyses.

## Extended Figure LEGENDS

**Extended Figure 1.** Examples of simple and complex SV patterns.

**Extended Figure 2.** Per-sample counts of complex (lower) and classified (upper) SV breakpoint junctions by histology group.

**Extended Figure 3.** Further examples of templated insertion chains, cycles and bridges.

Cancer-associated genes are shown in navy.

**Extended Figure 4.** Possible explanations for interlocked duplications.

(A) Two unphased fold-back inversions (through, for example, breakage-fusion-bridge events) could generate the structure.

(B) An extrachromomal (episomal) ring comprising the two segments linked by inverted rearrangements would recapitulate the rearrangements, but not the stable integer copy numbers generally observed.

(C) A series of unbalanced translocations between duplicated copies of the same chromosome is formally possible, but unlikely because of the close proximity of the rearrangements and stable background copy number of the chromosome.

(D-E) A tandem duplication, followed by inversion, followed by deletion could generate the structure, but few examples of the intermediate phase were observed.

(F) An example of such an event in a stomach cancer, occurring on the background of a single copy of the relevant chromosome arm. That this occurs within a single copy implies that the two inverted rearrangements must be phased, excluding the two fold-back inversion structure shown in (A).

**Extended Figure 5.** Further examples of local plus distant two-jump events.

These include (A) translocation with foldback, (B) translocation with insertion of a local inverted segment, and (C) translocation followed by tandem duplication spanning the translocation breakpoint junction (distinguishable from insertion cycles because of the uneven copy number either side).

**Extended Figure 6.** Size distribution of deletions and templated insertions.

(A) Size distribution of deletions per histology group, annotated with the number of deletions, number of samples, and the median deletion count per sample.

(B) Size distribution of deletions per sample, for a subset of individuals at deciles along the range of per-sample median sizes. Samples in the left panel had between 20 and 40 deletions; samples in the right panel had more than 40.

(C) Size distribution of segments of templated insertion, comparing events with one or multiple templated inserts.

(D) Comparison of the minimum and maximum templated insert size for multi-insert events.

(E) All events with three or more templated inserts, grouped by combination of insert sizes.

**Extended Figure 7.** Relationship of extended panel of genomic properties with SV categories.

Associations between genomic properties (rows) and classes of structural variation (columns). Each density curve represents the quantile distribution of the genomic property values at observed breakpoints compared to random genome positions, with stars indicating significant departure from uniform quantiles: FDR <0.01 *, <0.001 **, and <10^−6^ ***. Cells with significant property associations are shaded by the magnitude of the shift of the median observed quantile above (blue) or below (red) 0.5. The interpretation of each property from left to right is indicated in parentheses.

**Extended Figure 8.** Properties of SVs occurring in fragile sites.

(A) SV breakpoints in the most affected fragile sites: *FHIT, MACROD2* and *WWOX.* These are coloured by classification and vertically spaced by the distance to the next breakpoint in the cohort. If the two sides of a breakpoint junction are contained within the plotting window, they are joined by a curved line. The number of samples with a breakpoint in the plotting window is annotated top left.

(B) Number of deletions and tandem duplications (upper) and number of affected samples (lower) for the 18 fragile sites considered in this analysis.

(C) Size distribution of deletions and tandem duplications in fragile sites compared to the rest of the genome.

(D) Fragile site preference for 20 cancer histology groups as indicated by the proportion of samples harbouring a deletion in each of the 18 fragile sites considered here. The number of samples is indicated in parentheses.

**Extended Figure 9.** Distribution of SV signatures across tumour types. Per-sample SV footprint tallies considered for signatures analysis (upper) and estimated SV signature exposures (lower), as calculated by non-negative matrix factorization.

**Extended Figure 10.** Cancer genes disrupted by different categories of SV.

(A) SV breakpoints in the ERG locus, with examples of classic chromoplexy and complex chromoplexy generating the TMPRSS2-ERG fusion driver in prostate cancer.

(B) SV breakpoints in the PTEN locus, with an example of a templated insertion bridge disrupting PTEN in an ovarian cancer.

(C) SV breakpoints in the MDM2 locus, with an example of a complex event amplifying MDM2 in an osteosarcoma.

### SUPPLEMENTARY FIGURE LEGENDS

**Supplementary Figure 1.** Spearman correlations in library of genomic properties.

**Supplementary Figure 2.** Structural variation signatures estimated by the hierarchical Dirichlet process.

Shown are 90% credibility intervals. Here we show the estimated proportion of each event class assigned to each signature (rows sum to one).

**Supplementary Figure 3.** Structural variants classified by non-negative matrix factorisation.

**Supplementary Figure 4.** All templated insertion events affecting the *TERT* locus.

**Supplementary Figure 5.** *TERT* expression (log_10_ FPKM – UQ normalised) by SV status in liver cancer samples with available RNA.

**Supplementary Figure 6.** All templated insertion events affecting the *RB1* locus.

**Supplementary Figure 7.** *RB1* expression (log_10_ FPKM – UQ normalised) by SV status in breast cancer samples with available RNA.

